# Structural analysis of 23S rRNA methylating enzyme Cfr reveals RNA binding determinants for methylation regiospecificity and antibiotic resistance

**DOI:** 10.1101/2025.09.18.677109

**Authors:** Michael Fruci, Annia Rodríguez Hernández, Tatiana Skarina, Lou Ann Verellen, Kaitlyn Tsai, Johanna M. Virta, Danica Galonić Fujimori, Alexei Savchenko, Peter J Stogios

## Abstract

The 23S rRNA methylating enzyme Cfr, found in pathogens including *Staphylococcus aureus, Clostridium difficile, Escherichia coli* and *Klebsiella pneumoniae*, confers resistance to phenicols, lincosamides, oxazolidinones (including linezolid), pleuromutilins, and streptogramins A (the PhLOPS_A_ phenotype). Cfr catalyses methylation of the C8 position of the A2503 base in 23S rRNA; the recognition site of the above antibiotic classes. Along with RlmN housekeeping enzyme, Cfr can also promote methylation of the C2 position of the same base. The molecular and structural basis of Cfr’s dual substrate specificity is not known, which hinders our ability to design Cfr-targeting inhibitors necessary to curb the PhLOPS_A_ resistance. Here, we present the first crystal structure of Cfr and a detailed analysis of its possible interactions with rRNA. Using structure-guided mutagenesis, mass spectrometry analysis of *in-cellulo* 23S rRNA methylated species, and *in-cellulo* resistance studies, we identify the key amino acids essential for Cfr methylation and multi-drug resistance activity. In particular, we found that Cfr’s Q329 residue is important for C8-specific methylation. These data provide a framework for further studies of the biochemistry, structure and inhibition of this important resistance determinant.

## Introduction

Antimicrobial resistance (AMR) is recognized as a significant threat to modern medicine. The cause of AMR rise in clinically relevant microbial strains is multi-factorial: misuse and over-use of antibiotics in clinical, veterinary, agricultural settings^2^; pollution of the environment with antimicrobials and their metabolites or degradation products^3^; the inherent ability of bacteria to evolve resistance to chemical stresses^4^; the fact that most antibiotics are or are derived from natural products, which bacteria have been exposed to and evolved defenses against for eons^5,6^. Compounding the issue, bacteria have a propensity to exchange AMR genes (ARGs) on mobile genetic elements^7^. To tackle a global issue as complex as the AMR crisis, scientific insights must advance in numerous fields as well. This includes a more comprehensive understanding of the scope of possible resistance mechanisms and ARGs, their dissemination in the global environmental and clinical microbiomes, and detailed structural, biochemical and molecular data underlying the mechanistic basis of AMR-conferring proteins.

The ribosome is a common target for many different antibiotic classes, including tetracyclines, aminoglycosides, macrolides, phenicols, lincosamides, oxazolidinones, pleuromutilins, and streptogramins^8^. While specific interactions with the ribosome vary, a commonality among these drugs is their interaction with ribosomal RNA (rRNA), which corrupts essential processes in translation.

The PhLOPS_A_ antibiotics, which encompass phenicols, lincosamides, oxazolidinones (including linezolid), pleuromutilins, and streptogramins A, interact with the peptidyl transferase center (PTC) and 23S rRNA of the ribosome^8^. Multiple mechanisms of resistance to ribosome-targeting antimicrobials exist, including but not limited to chemical modification of the drug itself, which reduces drug affinity for its target (i.e. aminoglycoside-modifying enzymes^9^), ribosome-protecting proteins, including ABC-F family proteins^10^, which release antibiotics from the ribosome, expression of efflux pumps that eject antibiotics from the cell (e.g. Acr-Tol and Mex-Opr systems^11^, or chemical modification of the ribosome itself, typically by methylation.

rRNA methylation can be part of the housekeeping cell process in bacteria, providing a regulation mechanism for maturation and assembly of the ribosome, for modification of translation rate and efficiency, and for stress response (reviewed in ^12,13^). However, this reaction can also be “weaponized” to confer resistance to antibiotic drugs. For example, methylation of sites in 23S rRNA catalyzed by erythromycin resistance methyltransferase Erm and Radical S-adenosylmethionine (SAM) methylating enzyme Cfr^14^ confers antibiotic resistance. In these instances, methylation is thought to reduce the affinity of the antibiotics for binding to their target binding site(s). The *cfr* gene was discovered in 2000 as part of a plasmid in *Staphylococcus sciuri* that conferred *c*hloramphenicol and *f*lorfenicol *r*esistance (hence *cfr*) ^15,16^. It has since been discovered to confer PhLOPS_A_ resistance in many species^17^, including *S. aureus, Clostridioides spp*., *Enterococcus* spp., *Bacillus* spp., *Proteus vulgaris, Escherichia coli, Klebsiella pneumoniae*, and *Streptococcus suis*^17–22^. *Cfr* is associated with mobile genetic elements in most cases, indicating a selective advantage in its dissemination and increasing the importance of understanding its mechanism of action due to its mobility.

Cfr is a member of the Radical SAM Superfamily of enzymes (reviewed in ^23,24^); a large protein family supporting a diverse set of catalytic reactions. Cfr shares significant sequence similarity with RlmN (34% identity with the *E. coli* enzyme)^25^. RlmN catalyzes the methylation of the 23S rRNA’s A2503 base at the C2 position as well as the C2 position of A37 in tRNAs^25,27,28^ as part of large ribosome subunit assembly^26^. Notably, its activity does not confer PhLOPS_A_ resistance.

Expressed in the *E. coli rlmN* deletion strain and *in vitro*, Cfr was shown to catalyze methylation of A2503 at both the C2 and C8 positions. Cfr methylation on the C8 position conferred the PhLOPS_A_ phenotype^14,29^, consistent with the notion that C2 methylation is not responsible for resistance. We previously demonstrated that Cfr recognizes domain V of E. coli rRNA *in vitro*, with the 179 base region comprising 2447-2625 able to serve as a substrate for this enzyme^14^, while shorter rRNA fragments significantly compromise its methylation activity. Cryo-electron microscopy studies of Cfr-modified ribosomes identified that di-methylation of A2503 provided not only steric hindrance for binding of antibiotics, but also rearrangement of the local environment of the PTC, especially A2062, such that this disrupted the binding site for the antibiotics^30–32^.

The evolutionary origin of Cfr is thought to be either via gene duplication of the common ancestor of *rlmN* followed by diversification of the two enzymes towards C2-only and C2+C8 specificity, or via evolution of a Cfr-like family found in Firmicutes, including *Clostridium difficile*^33–36^. Therefore, a fundamental question remains: how is Cfr, while structurally and functionally similar to RlmN, also able to methylate the C8 position of A2503 to confer resistance? Previous analysis using the enzyme’s primary sequence analysis suggested that specific amino acids contribute to Cfr methylation activity, but this hypothesis was not validated^34^. Further understanding of the molecular activity of Cfr has been hampered by the lack of structural information on the Cfr enzyme. Such a knowledge gap also hindered the development of Cfr-specific inhibitors that would curb the resistance conferred by this enzyme.

In this study, we present the first crystal structure of the Cfr enzyme and a model of Cfr in complex with rRNA, leveraging the AlphaFold3 algorithm and experimentally defined structures of Cfr and 23S rRNA. We also tested the role of Cfr-specific residues in the enzyme’s active site by mutagenesis and expressing *cfr* variants in RlmN-null *E. coli*. This revealed multiple amino acids essential for Cfr methylation activity and resistance, and one residue in particular, that, when substituted for the equivalent RlmN-derived residue, resulted in a RlmN-like methylation pattern and abrogation of resistance.

## Results

### Crystal structure of the Cfr apo enzyme

To provide a molecular and structural framework of Cfr’s activity on two different sites on its substrate, we pursued structural characterization of this protein using X-ray protein crystallography. Accordingly, we recombinantly expressed and purified Cfr in the absence and in the presence of Fe^2+^, SAM cofactors and/or ribonucleotides. In addition to the wild-type Cfr (Cfr WT), we purified and attempted crystallization of the Cfr gain-of-function variants that we previously identified by a directed evolution approach^30^. Overexpression of Cfr in *E. coli* resulted in a bright red colored cell lysate, which we attributed to the overexpression of Cfr in complex with [4Fe-4S] cluster. This coloration was, however, lost during purification, probably due to the high sensitivity of this form of Cfr to oxygen exposure. Nonetheless, we obtained purified protein samples in case of Cfr WT as well as CfrV2 (N2K, S39G), CfrV3 (N2K, I26M, S39G) and CfrV4 (N2K, I26M, L68F). Unfortunately, we could not identify crystallization conditions for these protein samples.

To increase the probability of crystallization, we undertook Cfr sequence optimization using the PROSS server^37^. According to this server’s suggestions, we designed and tested for expression in *E. coli* 17 Cfr variants, which contained from 8 to 35 single amino acid substitutions, in addition to the gain-of-function mutations N2K, I26M, and L68F mentioned above^30^. Two of these Cfr variants selected based on their expression level in *E. coli*, were purified and screened for crystallization. One variant harbouring 18 individual amino acid substitutions (see the sequence in **Materials and Methods**), which we hereafter termed Cfr^PROSS-8,^ formed crystals that diffracted to 2.50 Å. Using diffraction data collected from this sample, we solved the Cfr^PROSS-8^ structure using Molecular Replacement (x-ray crystallographic statistics in **Supporting Information Table S1**), thus obtaining the first experimentally defined image of Cfr.

The crystal structure of Cfr^PROSS-8^ contained one polypeptide chain in the asymmetric unit corresponding to residues 1 to 329 of Cfr, in accordance with the design of our expression construct. Cfr protein adopted the canonical Radical SAM domain fold (InterPro IPR007197), consisting of a central 9-stranded β-sheet arrayed with 12 α-helices on the convex face of the β-sheet (**Figure 1**). The concave face of the β-sheet forms a cleft expected to contain the active site of the enzyme. However, the density for three regions in the Cfr structure, which correspond to the amino acids 111-125, 295-303 and 330-349, could not be resolved. In a previously determined crystal structure of RlmN, the regions equivalent to Cfr’s residues 111-125 and 295-303 contribute to the formation of the active site, and the region equivalent to Cfr 327-349 contains the conserved cysteine residue (RlmN C355, Cfr C338) essential for the catalytic activity.

**Figure 1.**
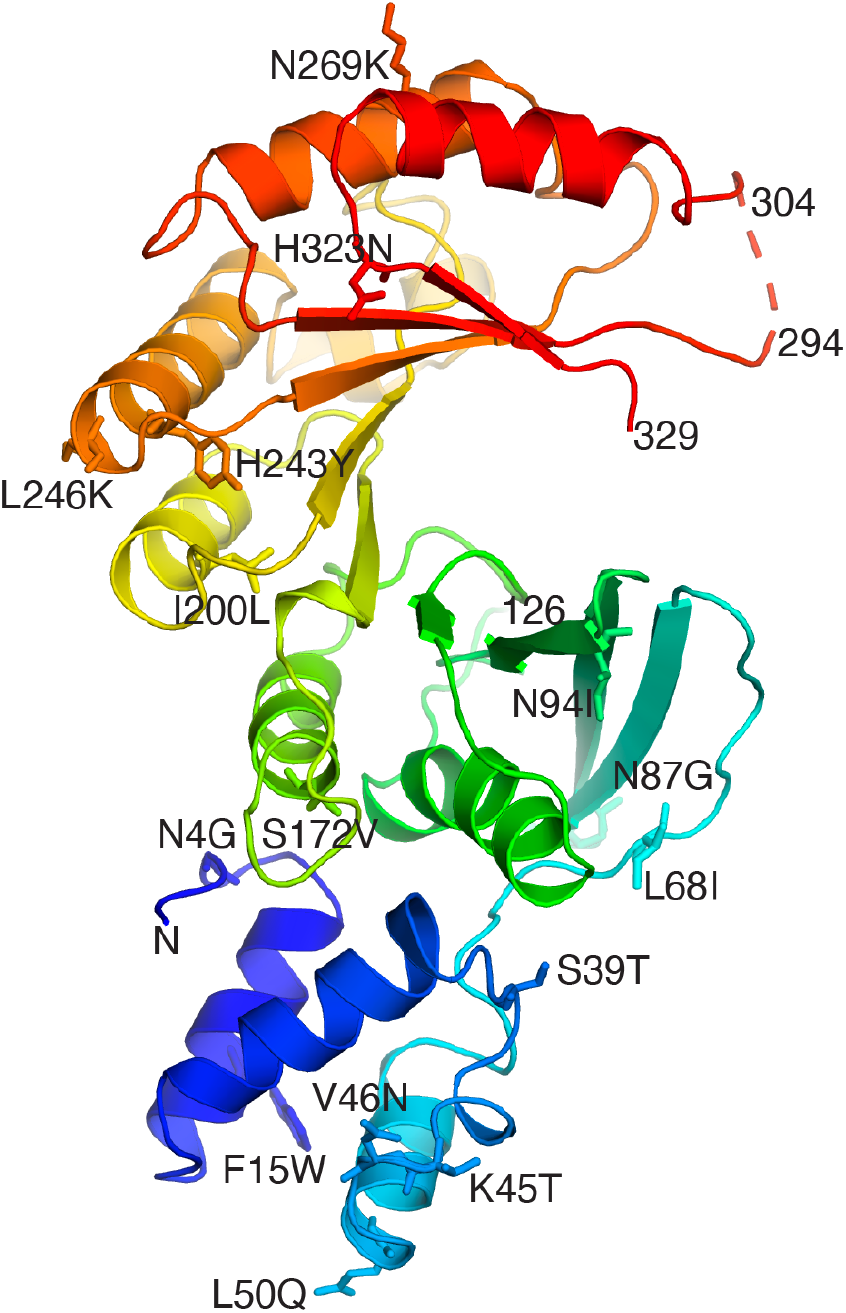
Crystal structure of Cfr^PROSS-8^. Labels indicate positions of N-terminus, C-terminally resolved residue (Q329), and PROSS-designed substitutions. Dashes indicate disordered positions.

We did not observe any additional electron density corresponding to the [4Fe-4S] cluster, SAM, or any other ligand and cofactors, suggesting that the obtained structure represents Cfr in the apo form. Accordingly, we hypothesized that the poor resolution of active site regions in the obtained structure is probably due to the lack of cofactors and ligands, resulting in increased flexibility of these regions of the enzyme. We were, nevertheless, able to unambiguously assign the density to sixteen residues (N4G, F15W, S39T, K45T, V46N, L50Q, L68I, K86D, N87G, N94I, S172V, I200L, H243Y, L246K, N269K, H323N) representing the introduced substitutions in Cfr^PROSS-8^ compared to the Cfr WT. These residues were distributed throughout the solvent-exposed surface of the enzyme but were distanced from the active site cleft (**Figure 1**). Three additional substitutions - N2K, I26M, L68F in Cfr^PROSS-8^, resulting in increased activity of Cfr^30^, were localized to the α-helical N-terminal lobe, also at a significant distance from the active site, suggesting that the activation effect of these mutations may be allosteric.

The Cfr^PROSS-8^ crystal structure superposes with RlmN from its complex with tRNA (PDB 5hr6^27^) with root-mean-square-deviation (RMSD) of 1.6 Å over 252 matching Cα atoms. Due to this significant structural similarity, we used the coordinates of the [4Fe-4S] cluster, a methionine and a 5’-deoxyadenosine (5’-dA) from the RlmN-tRNA complex to create a model of the Cfr *holo-*enzyme. The [4Fe-4S] cluster, the Met and 5’-dA were well accommodated into the Cfr active site using this strategy, and key interactions between Cfr and these ligands closely resembled those observed in the RlmN-tRNA complex, such as coordination of the [4Fe-4S] cluster by C112, C116 and C119, and the 5’-dA molecule was accommodated into a pocket comprised of F118, C119, M155, I290, C338 and G339 (residues all conserved in RlmN). We also generated a model of the Cfr *holo-*enzyme using Boltz-2^38^ at https://neurosnap.ai, and this closely resembled the model we generated by simply overlaying the Cfr^PROSS-8^ and RlmN crystal structures (not shown).

### Modelling of Cfr-23S rRNA complex

To gain insight into interactions between Cfr and its substrate, we modelled Cfr-RNA complex using the AlphaFold3 server (https://alphafoldserver.com). As the Cfr substrate submitted to modelling, we used the fragment of domain V (bases 2447-2635) of the *E. coli* 23S rRNA, previously shown to be sufficient to support the A2503 base methylation by Cfr^14^. The average predicted local distance difference test (pLDDT) score for the enzyme was 87.4, and 54.5 for the rRNA, respectively, indicating overall high and medium confidence for predicted positions of enzyme and rRNA in the model. At the MolProbity server (https://molprobity.biochem.duke.edu/index.php), the all-atom clashscore was 2.47, which was within the 99^th^ percentile, indicating very good model quality. We assessed the model generated by AlphaFold3 against the obtained Cfr^PROSS-8^ structure and the previously defined structure of RlmN in complex with tRNA (PDB 5hr6^27^). The model of 23S rRNA superimposed with RMSD 2.72 Å over 2947 atoms with the structure of the region comprising 2447-2583 bases of 23S rRNA from *E. coli* 70S ribosome (PDB 4v7d^39^) (**Supporting Information Figure S1**). Furthermore, the model suggested for Cfr in this complex was highly similar to the crystal structure of Cfr^PROSS-8^ (RMSD 1.17 Å over 284 matching Cα atoms). Based on this analysis, we deemed the suggested model highly confident and used it in further analysis.

According to the suggested model (**Figure 2**), the positively charged surface of Cfr that extends from the apex of the Radical SAM domain fold to its bottom and interacts with the rRNA. The A2503 base enters the active site that is itself positively charged (**Supporting Information Figure S2**). Notable positively charged residues from Cfr that interact with rRNA in this model include K27, K35, K98, R184, R185, K250, K280, K283, K315, K318, and stacking interactions include those between R184 and C2499 plus U2500, and between R250 and A2453 plus G2454 (**Supporting Information Figure S2**). A2503 is flipped out from the core of the RNA fold and is facing into the Cfr active site (**Figure 2A, 2B**). The model allowed to identify a number of amino acid sidechains approaching within 5 Å of the position of A2503, including M95, Y97, W101, S103, C105, D150, S151, S153, M155, R327, S328, Q329, F330, A337 and C338.

**Figure 2.**
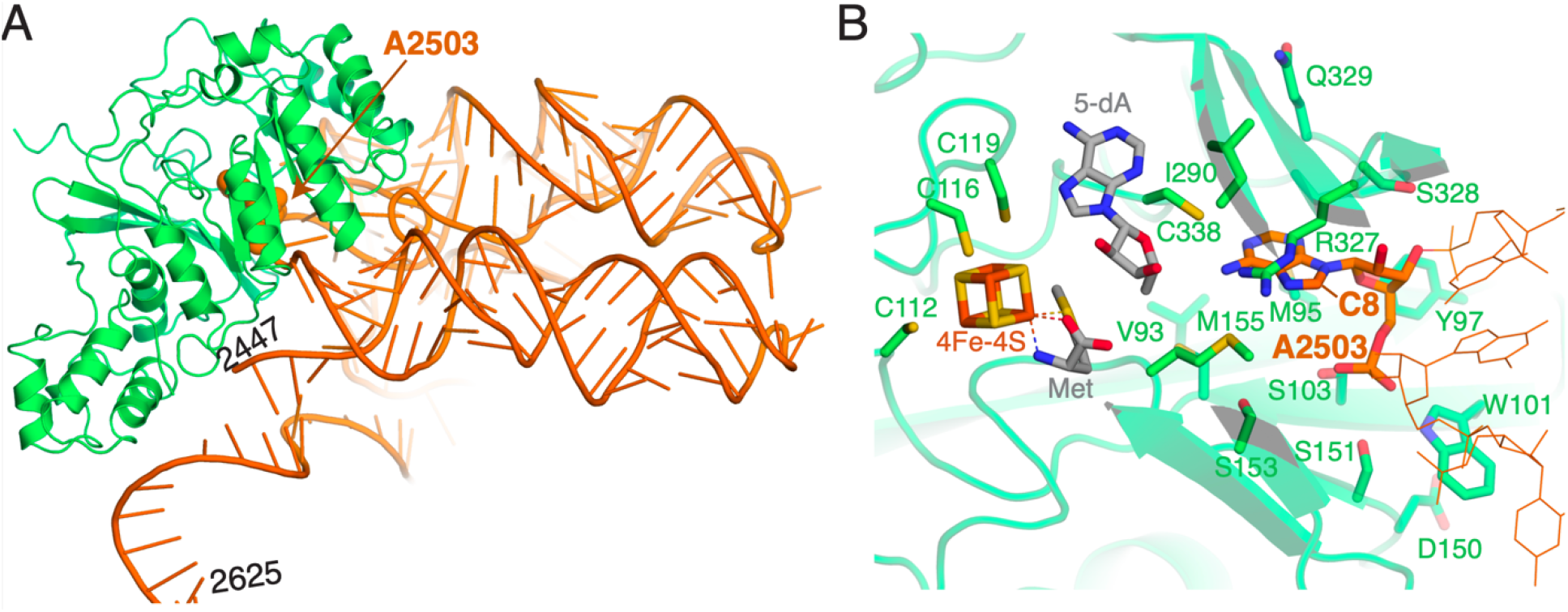
AlphaFold3-generated model of Cfr bound to its ligands and a fragment of *E. coli* 23S rRNA. A) Cfr shown in green cartoons, *E. coli* 23S rRNA shown in orange, with bases 2447, 2503 and 2625 labelled. B) Zoom of the Cfr active site showing locations of cofactors and A2503 (shown in thicker sticks and C8 labelled), remaining rRNA in thin lines. Amino acids labelled are those targeted for mutagenesis and/or key residues for interacting with co-factors.

Cfr in the model with rRNA are highly similar in structure to RlmN (RMSD 1.6-1.8 Å over 222-252 matching Cα atoms) (**Figure 3A**). Comparing the Cfr-rRNA and RlmN-tRNA models, both enzymes bound RNA on the equivalent face of the respective enzymes. Since this analysis confirmed significant structural similarity between RlmN and Cfr including the active site location, interactions with cofactors, and putative RNA binding surfaces, we hypothesized that the difference in methylation site specificity is dictated by diversification of specific active site residues in proximity to the cognate RNA base. Focusing on the residues within 5 Å of the position of A2503 in the Cfr-rRNA model, comparison of these 14 residues in Cfr with their equivalents in RlmN identified that 10 residues show significant variation, including: Cfr residues M95, Y97, W101, S103, D150, S151, S153, S328, Q329, F330 vs. RlmN residues I109, E111, R114, T116, T171, N172, V174, K345, T346, R347, respectively (**Figure 3B**). We thus hypothesized that these residues may be determinants of the C8 methylation specificity of Cfr as compared to the C2 specificity of RlmN.

**Figure 3.**
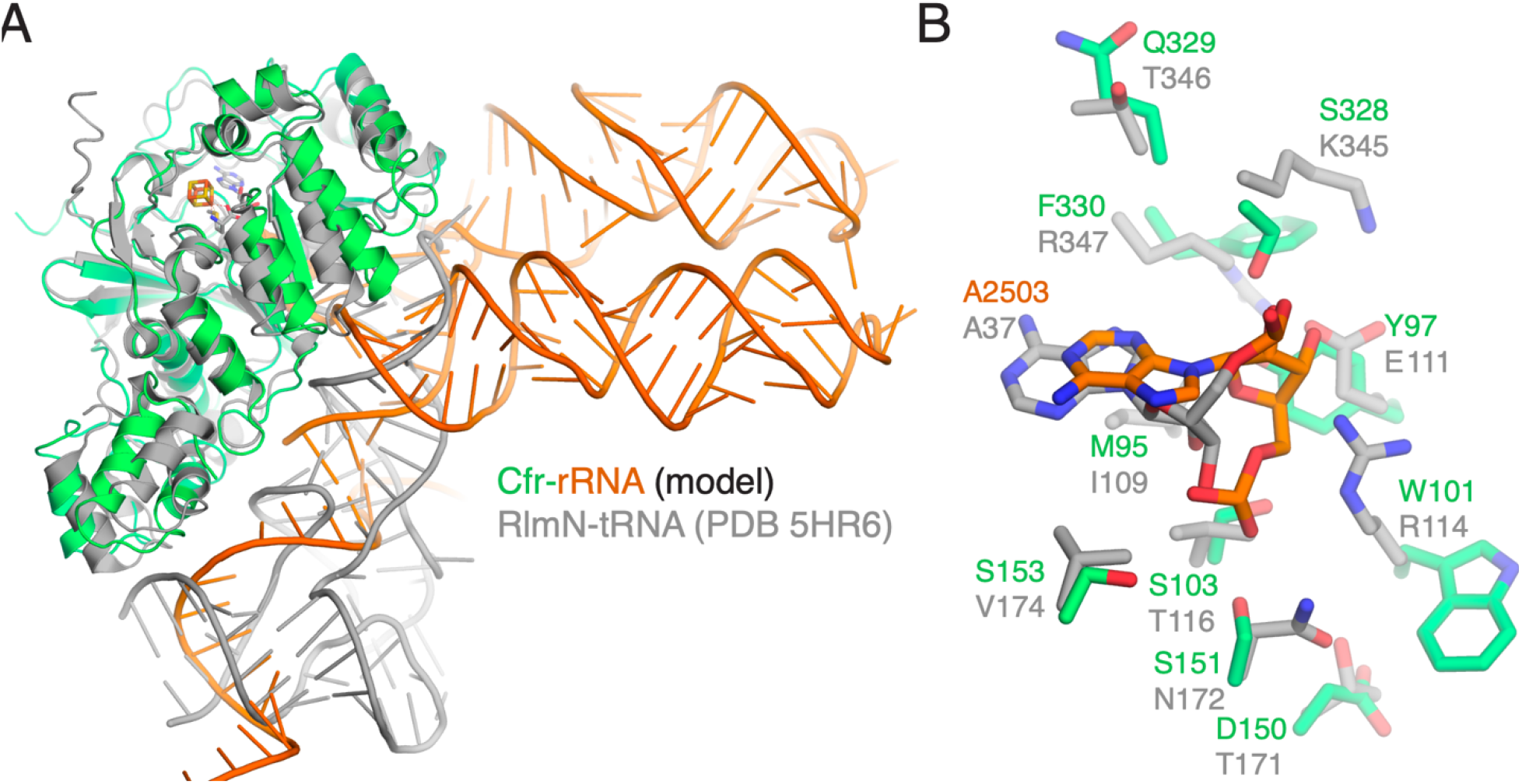
Comparison of the Cfr-rRNA AF model and the crystal structure of RlmN-tRNA complex. A) The Cfr-rRNA model is shown in green/orange, RlmN-tRNA (from PDB 5hr6^27^) is shown in grey. B) Zoom of active sites, with the same coloring scheme as panel A. Amino acids shown are those differing between Cfr and RlmN and targeted for mutagenesis in this study.

### Mutagenesis analysis of Cfr reveals key substrate-binding residues

Next, we tested our hypothesis about the role of specific amino acid residues in Cfr activity on the C8 position related to PhLOPS_A_ resistance by mutagenesis. We introduced individual substitution of selected residues into the *cfr* gene from *S. aureus*. These *cfr* variants were introduced on the pGDP2 expression plasmid into the hyperpermeable and efflux pump-deficient, PhLOPS_A_-sensitive strain of *E. coli* BW25113 Δ*bamB*Δ*tolC*^40^. Consistent with previous results, deletion of *bamB* and *tolC* conferred increased susceptibility to PhLOPS_A_ antimicrobials (**Table 1**)^40^ of this strain. This trend was reversed in the strain carrying plasmid pGDP2 expressing wild-type (WT) *cfr* gene. To confirm that the increased resistance is specific to PhLOPS_A_ and not 23S rRNA targeting antimicrobials in general, susceptibility to the non-PhLOPS_A_, 23S rRNA targeting antibiotic, erythromycin, was assessed. In agreement with Cfr-triggered resistance to PhLOPS_A_ antimicrobials, the strain carrying the pGDP2 plasmid expressing WT *cfr* did not alter susceptibility to erythromycin.

**Table 1.**
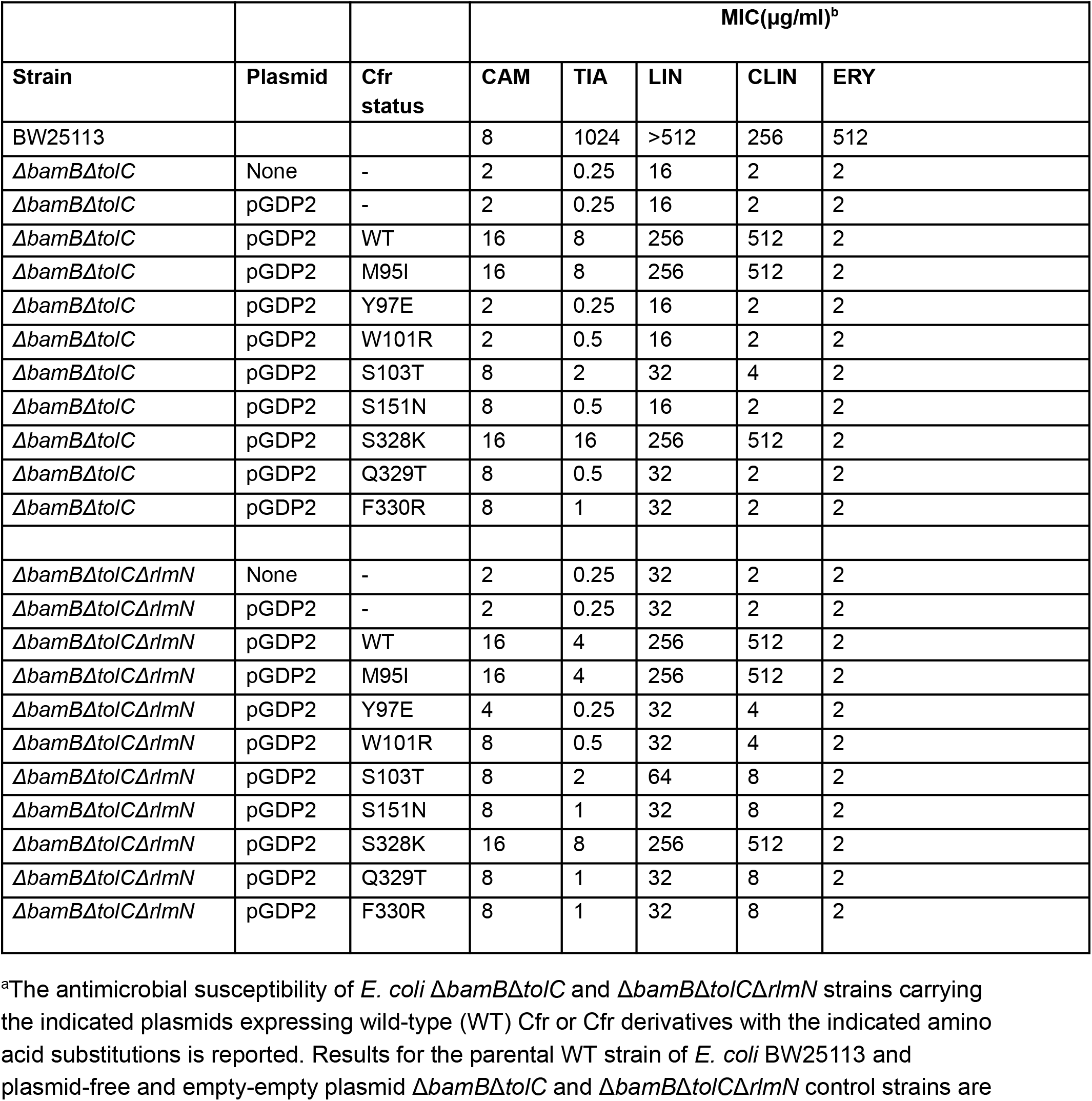
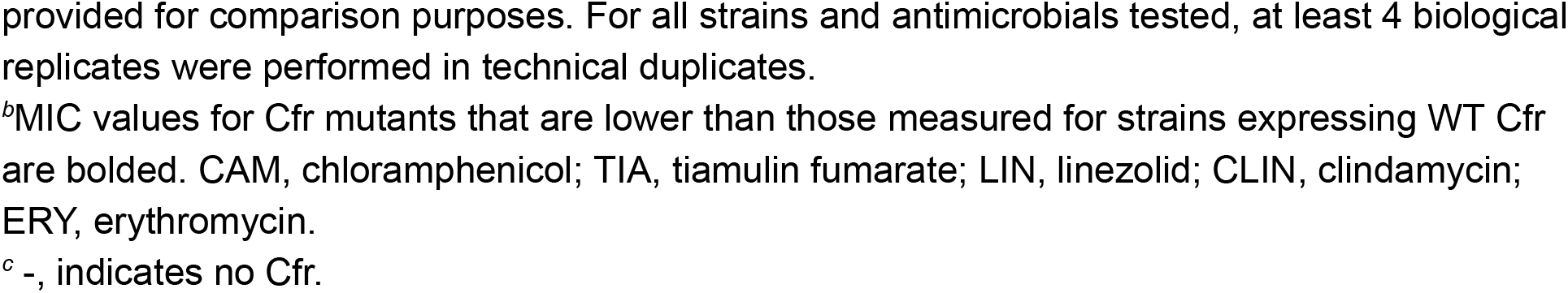
Impact of Cfr mutations on PhLOPS_A_ and erythromycin susceptibility of hyperpermeable strains of *E. coli*^*a*^.

| Strain | Plasmid | Cfr status | MIC(μg/ml)<sup>b</sup> |  |  |  |  |
| --- | --- | --- | --- | --- | --- | --- | --- |
|  |  |  | CAM | TIA | LIN | CLIN | ERY |
| BW25113 |  |  | 8 | 1024 | >512 | 256 | 512 |
| *ΔbamBΔtolC* | None | - | 2 | 0.25 | 16 | 2 | 2 |
| *ΔbamBΔtolC* | pGDP2 | - | 2 | 0.25 | 16 | 2 | 2 |
| *ΔbamBΔtolC* | pGDP2 | WT | 16 | 8 | 256 | 512 | 2 |
| *ΔbamBΔtolC* | pGDP2 | M95I | 16 | 8 | 256 | 512 | 2 |
| *ΔbamBΔtolC* | pGDP2 | Y97E | 2 | 0.25 | 16 | 2 | 2 |
| *ΔbamBΔtolC* | pGDP2 | W101R | 2 | 0.5 | 16 | 2 | 2 |
| *ΔbamBΔtolC* | pGDP2 | S103T | 8 | 2 | 32 | 4 | 2 |
| *ΔbamBΔtolC* | pGDP2 | S151N | 8 | 0.5 | 16 | 2 | 2 |
| *ΔbamBΔtolC* | pGDP2 | S328K | 16 | 16 | 256 | 512 | 2 |
| *ΔbamBΔtolC* | pGDP2 | Q329T | 8 | 0.5 | 32 | 2 | 2 |
| *ΔbamBΔtolC* | pGDP2 | F330R | 8 | 1 | 32 | 2 | 2 |
| *ΔbamBΔtolCΔrlmN* | None | - | 2 | 0.25 | 32 | 2 | 2 |
| *ΔbamBΔtolCΔrlmN* | pGDP2 | - | 2 | 0.25 | 32 | 2 | 2 |
| *ΔbamBΔtolCΔrlmN* | pGDP2 | WT | 16 | 4 | 256 | 512 | 2 |
| *ΔbamBΔtolCΔrlmN* | pGDP2 | M95I | 16 | 4 | 256 | 512 | 2 |
| *ΔbamBΔtolCΔrlmN* | pGDP2 | Y97E | 4 | 0.25 | 32 | 4 | 2 |
| *ΔbamBΔtolCΔrlmN* | pGDP2 | W101R | 8 | 0.5 | 32 | 4 | 2 |
| *ΔbamBΔtolCΔrlmN* | pGDP2 | S103T | 8 | 2 | 64 | 8 | 2 |
| *ΔbamBΔtolCΔrlmN* | pGDP2 | S151N | 8 | 1 | 32 | 8 | 2 |
| *ΔbamBΔtolCΔrlmN* | pGDP2 | S328K | 16 | 8 | 256 | 512 | 2 |
| *ΔbamBΔtolCΔrlmN* | pGDP2 | Q329T | 8 | 1 | 32 | 8 | 2 |
| *ΔbamBΔtolCΔrlmN* | pGDP2 | F330R | 8 | 1 | 32 | 8 | 2 |

<sup>a</sup>The antimicrobial susceptibility of *E. coli* *ΔbamBΔtolC* and *ΔbamBΔtolCΔrlmN* strains carrying the indicated plasmids expressing wild-type (WT) Cfr or Cfr derivatives with the indicated amino acid substitutions is reported. Results for the parental WT strain of *E. coli* BW25113 and plasmid-free and empty-empty plasmid *ΔbamBΔtolC* and *ΔbamBΔtolCΔrlmN* control strains are
provided for comparison purposes. For all strains and antimicrobials tested, at least 4 biological replicates were performed in technical duplicates.
<sup>b</sup>MIC values for Cfr mutants that are lower than those measured for strains expressing WT Cfr are bolded. CAM, chloramphenicol; TIA, tiamulin fumarate; LIN, linezolid; CLIN, clindamycin; ERY, erythromycin.
<sup>c</sup> -, indicates no Cfr.

Of note, expression of *cfr*^*PROSS-8*^ in *E. coli* failed to confer resistance to chloramphenicol (**Supporting Information Figure S3**), indicating that these introduced mutations had a negative effect on the activity of this enzyme. Given their locations away from the presumed active site of the enzyme, we hypothesized that these substitutions had an allosteric inhibition effect on Cfr.

Next, we assessed the antibiotic susceptibility of the same strain carrying a pGDP2::*cfr* plasmid encoding a variant with one of the following single amino acid substitutions: M95I, Y97E, W101R, S103T, S151N, S328K, Q329T and F330R. According to our model of Cfr-RNA complex, these amino acids localize in the predicted RNA-binding site. Individual mutations were designed to introduce a residue observed in RlmN at the equivalent position (**Figure 3B**). Most substitutions (i.e. Y97E, W101R, S103T, S151N, Q329T, R330R, **Table 1**) compromised resistance to all PhLOPS_A_ antimicrobials tested while not affecting the susceptibility to erythromycin. To rule out the variation in enzyme expression level, all strains were monitored for expression level and we did not observe any differences among WT and the variants **(Supporting Information Figure S4**). Only variants carrying M95I or S328K substitutions demonstrated the level of resistance comparable with that conferred by the Cfr WT (**Table 1**). Due to C2-A2503 methylation^25^ activity of RlmN, we next assessed the impact of the Cfr mutations in the absence of any endogenous A2503 methylation. To assess this, plasmid-borne Cfr WT and variants were introduced into a hyperpermeable, A2503 C2-methylation-deficient *E. coli* BW25113Δ*bamB*Δ*tolCΔrlmN* strain. Loss of endogenous A2503 methylation at C2 in the Δ*bamB*Δ*tolC* background did not alter the susceptibility of *E. coli* to PhLOPS_A_ antibiotics or erythromycin when compared to the Δ*bamB*Δ*tolC* strain. However, a modest 2-fold increase in resistance to linezolid was observed for the *rlmN* null strain. Plasmid-borne expression of Cfr WT in the *rlmN* null strain conferred a similar level of resistance to PhLOPS_A_ antibiotics as observed in the case of the *rlmN* positive strain expressing Cfr WT. As with the Δ*bamB*Δ*tolC* strain, expression of Cfr variants Y97E, W101R, S103T, S151N, Q329T, and F330R in the *rlmN*-null background compromised resistance to all PhLOPS_A_ antimicrobials tested, while Cfr variants carrying M95I or S328K substitutions conferred resistance compatible with that of Cfr WT. In several instances, we observed a two-to four-fold difference in the minimum inhibitory concentration (MIC) value between the *rlmN* null and the *rlmN* positive strains expressing plasmid-borne WT or Cfr variants. In general, clindamycin and linezolid MICs were higher for *rlmN* null background strains carrying Cfr inactivating mutations, compared to their *rlmN-*positive counterparts. In all instances, substitution mutations did not affect Cfr enzyme expression levels or erythromycin susceptibility in the *rlmN* null background (**Supporting Information Figure S4**). These data suggest that our structural analysis allowed the identification of key amino acids for the resistance functionality of Cfr.

### Cfr variants show altered m^8^A2503 methylation

Reductions of MIC values for the PhLOPS_A_ antibiotics caused by the introduction of specific amino acid substitutions into Cfr expressed in *E. coli* motivated an investigation into the ability of these mutants to catalyze methylation of A2503 *in cellulo. Cfr* mutants were cloned into the pZA vector and expressed in *E. coli* BW25113 Δ*bamB*Δ*tolC*. The *in vivo* 23S rRNA methylation activity was assessed using an oligonucleotide protection assay coupled to MALDI-TOF (see Methods). We observed no or weak signal for the double methylated m^2^m^8^AψG fragment of the 23S rRNA in case of Cfr variants carrying mutations which resulted in the lowest MIC values in our resistance assays - Y97E, W101R, and Q329T (**Figure 4A**). This contrasts with the expression of Cfr.WT, which resulted in a robust signal corresponding to the m^2^m^8^AψG rRNA fragment (**Figure 4A**). As expected, no m^2^m^8^AψG rRNA fragment was detected in the control strain carrying an empty vector. In line with resistance assay results, where the strain expressing Cfr S328K variant demonstrated MICs similar to the strain expressing the Cfr WT, the strain carrying this Cfr variant also demonstrated the highest intensity of the double methylated m^2^m^8^AψG fragment (**Figure 4A**). This method allowed us to correlate the relative methylation levels between the m^2^AψG and the m^2^m^8^AψG fragments to the observed MIC values for other variants, establishing intermediary double methylation activity for S103T, S151N, and F330R Cfr variants.

**Figure 4.**
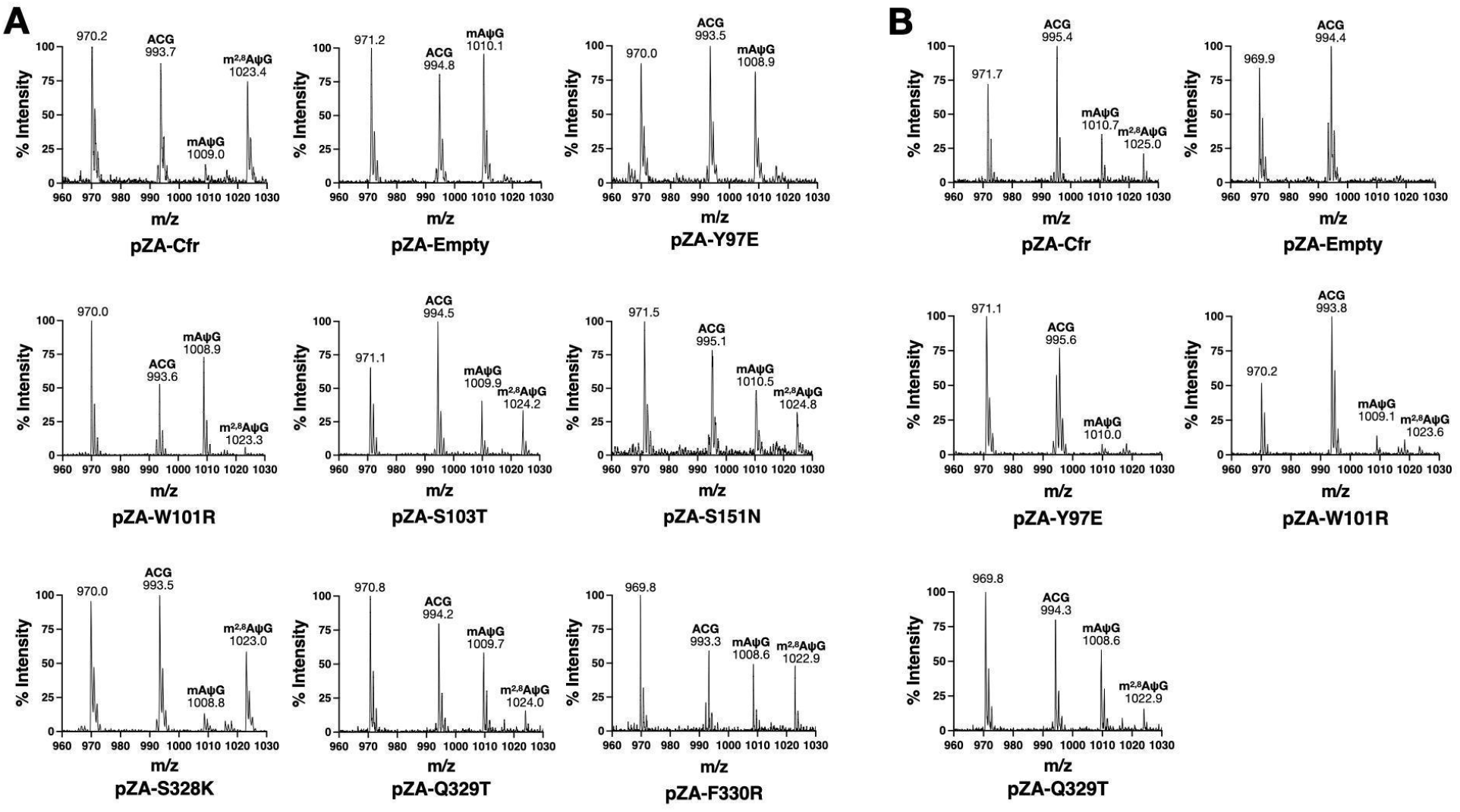
MALDI-TOF for A2503 containing rRNA fragments in two *E. coli* strains expressing Cfr and its variants. A) MALDI-TOF for rRNA fragments obtained from *E. coli* BW25113 Δ*bamB*Δ*tolC* cells expressing WT-Cfr, no-Cfr (empty vector), and Y97E, W101R, S103T, S151N, S328K, F330R, and Q329T Cfr single substitutions. C8 methylation activity of variants was evaluated by comparison of the mono-methylated mAψG and the double methylated m^2^m^8^AψG fragment. B) MALDI-TOF for rRNA fragments obtained from *E. coli* BW25113 Δ*bamB*Δ*tolC*Δ*rlmN* expressing WT-Cfr, empty vector, and Y97E, W101R, and Q329T Cfr variants.

While RlmN is considered a housekeeping enzyme with C2 methylation activity, in the absence of RlmN, Cfr can also catalyze C2 methylation in addition to C8 modification. To determine if the loss of the doubly methylated m^2^m^8^AψG fragment in the BW25113 Δ*bamB*Δ*tolC* strain was due to loss of catalytic activity or a change in methylation site specificity, we evaluated 23S rRNA methylation in the BW25113 Δ*bamB*Δ*tolC*Δ*rlmN* strain expressing the Cfr variants carrying one of Y97E, W101R and Q329T substitutions. As expected, the negative control strain transformed with an empty pZA vector displayed no signal for single and double methylation fragments (**Figure 4B**). When the same strain was transformed with the WT *cfr-*expressing plasmid, we observed the presence of both single and double methylation fragments (**Figure 4B**), similar to what was previously reported^40^; in the ratio of mono-to di-methylated product of 1.5:1. Given the very low intensity of mono-methylated product in the triple knockout strains expressing the Cfr Y97E or W101R variants (**Figure 4b)**, we hypothesized that the reduction in the second methylation event observed in the RlmN-expressing strain also expressing one of these two Cfr variants was likely due to loss of their catalytic activity. In contrast, expression of the Q329T variant led to the formation of both mono- and di-methylated products in the triple knockout strain (**Figure 4B**). Notably, the ratio of mono-to di-methylated product was higher in the strain expressing the Q329T variant (approximately 3-fold across biological replicates) (**Figure 4B**). Hence, we concluded that Q329T substitution shifts Cfr activity towards mono-methylation, therefore, suggesting that this residue is playing the key role in Cfr’s dual-methylation specificity.

## Discussion

Cfr mediates multi-drug resistance to PhLOPS_A_ antibiotics and is widely disseminated in bacterial pathogenic species. Cfr has been known to be highly related to RlmN in terms of sequence (both being members of the Radical SAM Superfamily) and function (23S rRNA methylation at A2503), and their corresponding genes likely share an evolutionary relationship through gene duplication^33^. Broader methylation specificity of Cfr relative to that of RlmN, which made the former enzyme a very important antibiotic resistance determinant, has raised an intriguing question of molecular mechanisms underpinning this unique activity by Cfr. However, the lack of structural information for Cfr has hindered our ability to address it. From the standpoint of enzyme evolution, it remains unknown how two closely related enzymes differ in their ability to methylate different positions on the same adenosine base in 23S rRNA, and the molecular determinants of recognition of the adenosine base that may contribute to this diversification.

In this study, we determined the first crystal structure of the Cfr enzyme, which relied on surface residue engineering via the PROSS server^37^. While the obtained crystal structure revealed the Cfr in an *apo* state lacking co-factors and substrate RNA, it provided a foundation for analysis of this enzyme’s molecular structure using advanced 3D modelling. Specifically, we leveraged the AlphaFold3 algorithm to generate a model of the *holo* Cfr-rRNA complex, which was validated using the *apo* crystal structure and enabled modelling of the enzyme in complex with its substrate.

Our validated model of Cfr-RNA complex provided the first insight into the role of specific amino acids involved in interactions with rRNA. Given the clear difference in methylation site preference on the A2503 base demonstrated by Cfr and RlmN enzymes, we reasoned that active site amino acids diversification is responsible for this phenomenon. Our subsequent resistance and *in vivo* methylation assays indicated that, among the investigated residues, Q239 likely contributes to the methylation site selectivity. One residue in the active site of Cfr (Q329), which, when substituted, results in heavily favoured mono-methylation of A2503 over di-methylation by the corresponding Cfr variant. The Cfr variant Q329T was also compromised for PhLOPS_A_ resistance, underscoring the importance of this particular amino acid.

Most of the individual substitutions of the amino acids in the Cfr active site that show diversification compared to the corresponding residues in RlmN resulted in loss-of-function effects for Cfr activity, preventing dissection of their roles in A2503 methylation specificity. Thus, further work is required to identify any additional residues that may be contributing to this Cfr activity.

The exact position of the introduced methyl group in mAψG species resulting from the activity of the Cfr Q329T variant cannot be determined from the assays described in this study. However, the formation of di-methylated species suggests that this Cfr variant, in analogy to the Cfr WT, has likely retained the ability to methylate both the C2 and C8 carbon atoms of A2503. The ability of the Cfr Q329T variant to methylate both the C2 and C8 positions of A2503 is in agreement with our earlier observations that the Cfr WT can also support the methylation of both amidine carbons of the substrate^14^. According to our model of Cfr, the Q329 residue is positioned at the periphery of this enzyme’s active site, and thus understanding its specific interactions with rRNA requires further structural and biochemical analysis of the Cfr-rRNA complex. Q329 is highly conserved in Cfr orthologs^34^, while RlmN orthologs typically feature a glutamate or threonine residue at a similar position. Many of the other amino acid positions that were interrogated by mutagenesis in this study are also highly conserved in Cfr orthologs, including Y97, W101, S103, S151, S153, Q329, and F330, while they are diversified in RlmN^34^.

Our comparative analysis of the Cfr-23S rRNA model with the crystal structure of the homolog RlmN bound to tRNA (PDB 5hr6) shows the RNA substrate positioned inside the active site cleft in the opposite orientation in the case of these two enzymes: 5’-3’ for RlmN and 3’-5’ for Cfr. This may or may not reflect the actual binding direction of the RNA in Cfr’s active site; however, if experimentally confirmed, such a difference in substrate binding could also affect positioning of the A2503 base in the catalytic cleft, favouring Cfr’s methyl transfer to the C8 over the C2 carbon. Therefore, substrate directionality might also play a part in the carbon specificity of these two enzymes.

The crystal structure of RlmN in complex with tRNA shows the C2 positioned towards the 5’-dA methylation byproduct^27^; we expect that Cfr can position the C8 site of A2503 for methylation, through interactions with Q329 and other residues in the active site. More extensive mutagenesis probing of Cfr’s active site, targeting positions along the putative rRNA binding surface not covered by the current study, can expand our knowledge on how Cfr recognizes rRNA. Additionally, expanding the structural, molecular and mutagenesis studies to Cfr orthologs, including those found in Bacillales^34^ and Clostridiales^41^, could identify additional important positions in the Cfr sequence for methylation specificity. Finally, the structural and molecular framework we provide in this study will be important for the design of Cfr inhibitors towards the development of adjuvant compounds capable of reversing Cfr-driven PhLOPS_A_ resistance^40^.

## Supporting Information

Evaluation of chloramphenicol resistance conferred by Cfr^PROSS-8^, Comparison of 23S rRNA coordinates from the Cfr-rRNA model and crystal structure of *E. coli* ribosome, Surface electrostatic charge of Cfr (from Cfr-rRNA AlphaFold3 model), Validation of expression of Cfr variants in *E. coli BW25113* Δ*bamB*Δ*tolC* and Δ*bamB*Δ*tolC*Δ*rlmN*, X-ray crystallographic statistics for Cfr^PROSS-8^, *E. coli* strains, plasmids and oligonucleotides used.

## Acknowledgements

We thank Rosa Di Leo for assistance with cloning. We thank Youngchang Kim of the Structural Biology Center, Advanced Photon Source, Argonne National Laboratory, for X-ray diffraction data collection and/or data processing. We thank Gerry Wright for providing plasmids pGDP2 and the strains *E. coli* Δ*bamB*Δ*tolC* and *E. coli* Δ*bamB*Δ*tolC rlmN::kan*. Research at AAFC was supported through AAFC A-base (J-002363) and the Government of Canada Shared Priority Project, Genomics Research and Development Initiative, project GRDI-AMR-One Health (J-002863). This work has been funded in whole or in part with U.S. Federal funds from the National Institute of Allergy and Infectious Diseases, National Institutes of Health, Department of Health and Human Services, under Contract Nos. HHSN272201700060C and 75N93022C00035 (Center for Structural Biology of Infectious Diseases, CSBID) to A.S and NIAID (R01 AI137270 to DGF). This research also used resources of the Advanced Photon Source, a U.S. Department of Energy (DOE) Office of Science User Facility operated for the DOE Office of Science by Argonne National Laboratory under Contract No. DE-AC02-06CH11357. The content is solely the responsibility of the authors and does not necessarily represent the official views of the National Institutes of Health. This manuscript is the result of funding in whole or in part by the National Institutes of Health (NIH). It is subject to the NIH Public Access Policy. Through acceptance of this federal funding, NIH has been given a right to make this manuscript publicly available in PubMed Central upon the Official Date of Publication, as defined by NIH.

## Abbreviations Used

4Fe-4S: 4 iron-4 sulfur cluster
5’-dA: 5’-deoxyadenosine
AMR: Antimicrobial resistance
ARG: Antimicrobial resistance gene
MIC: minimum inhibitory concentration
pLDDT: predicted local distance difference test
PhLOPS_A_: phenotype of resistance to phenicols, lincosamides, oxazolidinones, pleuromutilins, and streptogramins A
PTC: Peptidyl transferase center RMSD: root-mean-square-deviation
SAM: S-adenosylmethionine

## Materials and Methods

### Bacterial strains, plasmids and growth conditions

The bacterial strains and plasmids used in this study are described in **Supplemental Data Tables 1 and 2**, and oligonucleotide primers in **Supplemental Data Tables 3**. Bacterial cells were cultured in Luria-Bertani broth (LB-broth) and on Luria-Bertani agar (LB-agar) at 37°C, unless otherwise stated. In *E. coli*, plasmid pGDP2 and its derivatives were maintained with 25 µg/ml kanamycin, and pCP20 was maintained with ampicillin 100 µg/ml at 30°C.

For mutagenesis in the pZA vector, standard mutagenesis was used to introduce single mutation sites in the *cfr* gene for Y93E, W101R, S151N, S103T, S328K, F330R, and Q329T using the oligonucleotide sequences in **Supplementary Data Table 3**. Each mutant was confirmed by whole plasmid sequencing performed by Plasmidsaurus (South San Francisco, CA, USA) using Oxford Nanopore Technology with custom analysis and annotation. Once sequences were confirmed, plasmids were transformed into corresponding *E. coli* strains for expression. For pGDP2 expressing WT and variant *cfr* genes, constructs were designed *in silico*, synthesized and cloned into pGDP2 at Twist Bioscience (South San Francisco, CA, USA) and subsequently transformed into *E. coli* strains.

### Genetic Manipulation

To obtain a markerless deletion of the *rlmN* gene in the *E. coli* Δ*bamB*Δ*tolC*Δ*rlmN*::kan^R^ strain, plasmid pCP20 expressing FLP recombinase was introduced into this strain using a similar protocol as previously described^44^. *E. coli* cells were made competent by the CaCl_2_ method and subsequently transformed with pCP20. Transformants were allowed to recover for 2 hours at 30°C, and 20 µl of the culture was plated on LB agar plates containing 100 µg/ml of ampicillin (LB-AMP). Plates were incubated at 30°C for approximately 48 hours. Single colonies were then selected from the LB-AMP plates and inoculated into LB broth and incubated overnight at 43°C to select for loss of pCP20. Overnight cultures were then serially diluted and plated on LB plates and incubated for 24 hours at 30°C. To confirm loss of both pCP20 and the kanamycin resistance cassette, colonies were patched onto LB+Kan, LB+Amp and L-agar plates. The LB and LB+Kan plates were grown at 37°C, while the LB+AMP plates were grown at 30°C. *E. coli* Δ*bamB*Δ*tolC*Δ*rlmN* isolates that were sensitive to both kanamycin and ampicillin were selected and archived until further analysis.

### Engineering of Cfr for crystallization

The sequence of CfrV3^30^ and a model of Cfr generated using I-TASSER^45^ were submitted to the PROSS server^37^, which returned multiple variants of Cfr harboring between 8 and 40 amino acid substitutions. The designs were compared to filter only those substitutions to all designs, then further filtered to eliminate positions within the active site or possibly forming interactions with RNA, as suggested by superposition with the structure of the RlmN-tRNA complex^27^. This produced a list of 17 variants, all of which were purchased as synthesized genes from Twist Biosciences, then subcloned into pMCSG53. IPTG-inducible expression in *E. coli* BL21(DE3)-Gold cells was evaluated at a small scale (50 mL cultures) and via SDS-PAGE. Two variants, including two variants we named Cfr^PROSS-7^ (harboring substitutions N2K, I26M, L68F, K86D, N87G, T92A, N94I, K128R, R164P, S172V, I200L, S215A, Y217N, H243Y, N269K, V272A, S273K, V310I) and Cfr^PROSS-8^ (N2K, N4G, F15W, I26M, S39T, K45T, V46N, L50Q, L68I, K86D, N87G, N94I, S172V, I200L, H243Y, L246K, N269K, H323N) showed expression in the soluble fraction and were chosen for large scale purification.

### Protein purification, crystallization, x-ray diffraction and structure solution

*E. coli* BL21(DE3)-Gold competent cells were transformed with the pMCSG53 expression plasmid harboring *cfrPROSS-7* or *cfrPROSS-8* and purified according to previously published methods^46^. Both proteins were colorless throughout the purification, indicative of loss of the 4Fe-4S cluster. The final proteins (His_6_-tag cleaved) were dialyzed with 2 × 2 L dilution cycles in 10 mM HEPES at pH 7.5 with 300 mM NaCl, 0.5 mM TCEP. Purity was analyzed by SDS-PAGE and mass spectrometry. The proteins were concentrated using a Vivaspin concentrator (GE Healthcare) and passed through a 0.2 μm Ultrafree-MC centrifugal filter (Millipore) before storing in aliquots at −80 °C. Crystallization was performed at a room temperature of 21 °C by the sitting drop vapor diffusion method with 0.6 μL protein and/or protein: ligand substrate mix plus 0.6 μL reservoir solution, using a STP Labtech Mosquito robot. Crystals of Cfr^PROSS-8^ grew in drops with reservoir solution 40% v/v PEG400, 0.1 M HEPES pH 7.5, 7 mM β-mercaptoethanol, 0.1 M NDSB211. The crystal was cryoprotected by transferring to paratone oil, then flash frozen in liquid nitrogen. X-ray diffraction data at 100 K were collected at the beamline 19-ID, Structural Biology Center, Advanced Photon Source, Argonne National Laboratory, Argonne, IL, USA (wavelength 0.978 Å). All diffraction data were processed using HKL3000^47^. Molecular Replacement was utilized to solve the structure using the structure of the radical SAM methyltransferase from *Sphaerobacter thermophilus* (PDB 6FZ6) as the search model, with the CCP4 online server MrBump^48^. Model building and refinement were performed using Phenix.autobuild, Phenix.refine, and Coot^49,50^. Geometry was evaluated using Phenix.molprobity, and the wwPDB validation server. Atomic coordinates are available at the Protein Data Bank with accession code 9P0P. Production of figures and analysis was performed using The PyMOL Molecular Graphics System, Version 2.4.0, Schrödinger, LLC.

### Modelling of Cfr-23S rRNA complex

The DeepMind AlphaFold3 server (https://alphafoldserver.com) was used to generate a model with the sequence of the *E. coli* BW25113 23S rRNA (NCBI accession CP009273.1), using the region of rRNA (bases 2218-2615), which includes the region (2447-2625) previously identified as representing a good substrate for Cfr-catalyzed A2503 di-methylation^14^. Most returned models did not show significant buried surface area between Cfr and the rRNA, or interaction with the Cfr active site, except the model with bases 2447-2635, which was selected for further analysis. Coordinates of [4Fe-4S] cluster, a methionine and a 5’-deoxyadenosine (5’-dA) were added to the model through superposition with the structure of the RlmN-tRNA complex (PDB 5hr6^27^). The Cfr-23S rRNA model was energy minimized using the YASARA energy minimization server (https://www.yasara.org/minimizationserver.htm) and geometry was evaluated by the MolProbity server (https://molprobity.biochem.duke.edu/index.php).

### Cfr mutant expression and RNA recovery

The pZA vectors with the wild type sequence of *S. aureus* Cfr, and the Y93E, W101R, S151N, S103T, S328K, F330R, and Q329T mutants, as well as an empty pZA vector, were transformed into *E. coli* BW25113 Δ*bamB*Δ*tolC*, wild type, empty vector, Y93E, W101R, S328K, and Q329T were also transformed into the BW25113 Δ*bamB*Δ*tolC*Δ*rlmN::kan* strain. *E. coli* BW25113 Δ*bamB*Δ*tolC* transformants were plated in LB-agar with 100 μg/mL ampicillin, and *E. coli* BW25113 Δ*bamB*Δ*tolC*Δ*rlmN* on LB-agar with 100 μg/mL ampicillin and 50 μg/mL kanamycin to maintain the selective pressure of the aminoglycoside phosphotransferase conferring kanamycin resistance used as the replacement cassette for RlmN^40^. A single colony was selected from each plate, and 10 mL of overnight cultures with corresponding antibiotics were grown for 18 h at 37°C while shaking. The next day, 20 mL of fresh LB with antibiotics and 30 ng/mL anhydrotetracycline hydrochloride (AHT) were inoculated with 200 μL of the overnight culture. Cultures were grown at 37°C for 3-4 h until OD600nm reached 0.53-0.91, then cells were chilled on ice and centrifuged at 4°C for 10 min to remove media. Total RNA was purified using the RNeasy Midi Kit from Qiagen (Toronto, Ontario, Canada) by the manufacturer’s protocol. The average concentration of each RNA sample was obtained using a NanoDrop One^C^ instrument (Madison, WI, USA) by measuring the neat and a 1/10 dilution; an A260/A280 ratio of ≥ 2.0 was considered for quality. The RNA was flash frozen and stored at -20°C until needed.

### Total RNA digestion, isolation of A2503 containing fragments, and MALDI-TOF

Once total RNA was obtained, it was annealed to the C2480-C2520 DNA oligo complementary to 41 nucleotides in the *E. coli* 23S rRNA containing the A2503 nucleotide (**Supplementary Table 3**). The materials were digested and treated following the exact method previously published^40^. Briefly, after Mung Bean and RNase A nucleases treatment, fragments were precipitated, recovered by centrifugation, air dried, and resuspended in 24 μL of H_2_O:formamide: RNA loading buffer in a 1:2:2 ratio. To separate the rRNA fragments from the DNA oligo, samples were loaded in an 8 M urea, 13% polyacrylamide 1X TBE mini gel, run for 2 h at 100 V and developed by SYBR Gold stain. Gels were visualized in a BioRad ChemiDoc MP Imaging System, and the single rRNA fragment band was sliced in a Benchtop 2UV Transilluminator at 302 nm. Gel pieces were extracted in 1X TE buffer by heat and freeze cycles, and gel was removed with Spin X-centrifuge tubes (Costar 8160). rRNA was precipitated overnight in the presence of 1 μL glycoblue (Invitrogen AM9515), recovered by centrifugation, resuspended in an 18 μL solution of 0.14 M 3-Hydroxypicolinic acid (3-HPA) (Thermo Scientific 241050050) and 500 U of T1 nuclease (Thermo Scientific 01221951), and incubated for 3 hours at 37°C. After digestion, 0.12 M HCl was added, and the mix was incubated at ambient temperature for 30 min. Finally, fragments were flash frozen, lyophilized, and resuspended in 5 μL of UltraPure water (Invitrogen 10977-015). 2 μL of this suspension was sandwiched between 2 μL of freshly made 0.5 M 3-HPA on an MS 2mm 384 Well Sample Plate (Shimadzu, TO-454R00) and dried in a desiccator for 15 min. Data were acquired in a Shimadzu AXIMA Performance instrument, analyzed using the Shimadzu MALDI-MS software, and re-plotted using GraphPad Prism 10.2.3 software. Three to five technical repetitions were collected for each of the two biological repetitions tested in this study.

### Antibiotic Susceptibility Testing

The susceptibility of *E. coli* strains transformed with pGDP2 to clindamycin, tiamulin fumarate, linezolid, chloramphenicol and erythromycin was assessed using the 2-fold serial dilution technique in a 96-well microtiter plate as previously described^51^. Trailing growth was observed for the Δ*bamB*Δ*tolC and* Δ*bamB*Δ*tolC*Δ*rlmN E. coli* strains for all antibiotics tested, which made end-point determination difficult. As recommended by CLSI (CLSI. Performance standards for antimicrobial susceptibility testing. 30th ed. CLSI supplement M100. Wayne, PA: Clinical and Laboratory Standards Institute; 2020), the MICs for these antibiotics were defined as the lowest concentration where trailing began; tiny buttons of growth were ignored (i.e., the first well in which growth was significantly reduced). For clindamycin Δ*bamB*Δt*olC*Δ*rlmN* and Δ*bamB*Δ*tolC*, growth appeared to decrease with increasing clindamycin concentrations; however, a modest rebound in growth would occur at higher drug concentrations. In these instances, the MIC was read as the first well in which growth was significantly reduced.

### Whole-cell protein extracts and western immunoblotting

*E. coli* whole-cell protein extracts and western immunoblotting were performed similarly as previously outlined^46^. Blots were first probed with a primary antibody against -E. coli RNA polymerase β subunit (BioLegend, 1:5000) and a secondary HRP-conjugated goat anti-mouse IgG antibody (Abcam, 1:5000). Once imaged, primary and secondary antibodies were stripped from the blots with the Thermo Scientific Restore PLUS Western Blot Stripping Buffer according to a protocol outlined by the manufacturer. Following stripping, blots were reprobed with the primary anti-flag M2 (Sigma Aldrich, 1: 5000) and secondary HRP-conjugated polyclonal goat anti-mouse IgG H&L antibodies (Abcam, 1:5000). Blots were imaged using a Uvitec Alliance Q9 Advanced Chemiluminescence Imager (UVITEC Cambridge, Cambridge, England, UK).

## References

1. Naghavi, M. et al. Global burden of bacterial antimicrobial resistance 1990–2021: a systematic analysis with forecasts to 2050. The Lancet 0, (2024).

2. Hernando-Amado, S., Coque, T. M., Baquero, F. & Martínez, J. L. Defining and combating antibiotic resistance from One Health and Global Health perspectives. Nat. Microbiol. 4, 1432–1442 (2019).

3. Larsson, D. G. J. & Flach, C. F. Antibiotic resistance in the environment. Nat. Rev. Microbiol. 20, 257–269 (2022).

4. Baquero, F. et al. Evolutionary Pathways and Trajectories in Antibiotic Resistance. Clin. Microbiol. Rev. 34, e0005019 (2021).

5. Agudo, R. & Reche, M. P. Revealing antibiotic resistance’s ancient roots: insights from pristine ecosystems. Front. Microbiol. 15, 1445155 (2024).

6. Perry, J., Waglechner, N. & Wright, G. The prehistory of antibiotic resistance. Cold Spring Harb. Perspect. Med. 6, (2016).

7. Partridge, S. R., Kwong, S. M., Firth, N. & Jensen, S. O. Mobile genetic elements associated with antimicrobial resistance. Clin. Microbiol. Rev. 31, 1–61 (2018).

8. Lin, J., Zhou, D., Steitz, T. A., Polikanov, Y. S. & Gagnon, M. G. Ribosome-Targeting Antibiotics: Modes of Action, Mechanisms of Resistance, and Implications for Drug Design. Annu. Rev. Biochem. 87, 451–478 (2018).

9. Ramirez, M. S. & Tolmasky, M. E. Aminoglycoside modifying enzymes. Drug Resist. Updat. Rev. Comment. Antimicrob. Anticancer Chemother. 13, 151–171 (2010).

10. Sharkey, L. & O\textquoterightNeill, A. J. The antibiotic resistance ABC-F proteins: bringing target protection into the limelight. ACS Infect. Dis. acsinfecdis.7b00251 (2018).

11. Li, X.-Z., Plésiat, P. & Nikaido, H. The challenge of efflux-mediated antibiotic resistance in Gram-negative bacteria. Clin. Microbiol. Rev. 28, 337–418 (2015).

12. Pletnev, P. et al. Comprehensive Functional Analysis of Escherichia coli Ribosomal RNA Methyltransferases. Front. Genet. 11, 97 (2020).

13. Sergiev, P. V., Aleksashin, N. A., Chugunova, A. A., Polikanov, Y. S. & Dontsova, O. A. Structural and evolutionary insights into ribosomal RNA methylation. Nat. Chem. Biol. 14, 226–235 (2018).

14. Yan, F. et al. RlmN and Cfr are radical SAM enzymes involved in methylation of ribosomal RNA. J. Am. Chem. Soc. 132, 3953–3964 (2010).

15. Schwarz, S., Werckenthin, C. & Kehrenberg, C. Identification of a Plasmid-Borne Chloramphenicol-Florfenicol Resistance Gene in Staphylococcus sciuri. Antimicrob. Agents Chemother. 44, 2530–2533 (2000).

16. Long, K. S., Poehlsgaard, J., Kehrenberg, C., Schwarz, S. & Vester, B. The Cfr rRNA methyltransferase confers resistance to Phenicols, Lincosamides, Oxazolidinones, Pleuromutilins, and Streptogramin A antibiotics. Antimicrob. Agents Chemother. 50, 2500–2505 (2006).

17. Vester, B. The cfr and cfr-like multiple resistance genes. Res. Microbiol. 169, 61–66 (2018).

18. Yang, W., Chen, T., Zhou, Q. & Xu, J. Resistance to linezolid in Staphylococcus aureus by mutation, modification, and acquisition of genes. J. Antibiot. (Tokyo) 78, 4–13 (2025).

19. Sadowy, E. Linezolid resistance genes and genetic elements enhancing their dissemination in enterococci and streptococci. Plasmid 99, 89–98 (2018).

20. Sun, M. et al. Emergence of Multidrug-Resistant Proteus mirabilis Harboring both bla NDM-1 and cfr genes. Biomed. Environ. Sci. BES 37, 1208–1212 (2024).

21. Tang, Y. et al. Characterization of three porcine Escherichia coli isolates co-harbouring tet(X4) and cfr. J. Antimicrob. Chemother. 76, 263–264 (2021).

22. Feng, Y. et al. Identification and evolution of novel cfr-carrying plasmids in XDR Klebsiella pneumoniae strains from a chicken farm. Microbiol. Spectr. 13, e0262824 (2025).

23. Fujimori, D. G. Radical SAM-mediated methylation reactions. Curr. Opin. Chem. Biol. 17, 597–604 (2013).

24. Hoffman, B. M., Broderick, W. E. & Broderick, J. B. Mechanism of Radical Initiation in the Radical SAM Enzyme Superfamily. Annu. Rev. Biochem. 92, 333–349 (2023).

25. Toh, S.-M., Xiong, L., Bae, T. & Mankin, A. S. The methyltransferase YfgB/RlmN is responsible for modification of adenosine 2503 in 23S rRNA. RNA N. Y. N 14, 98–106 (2008).

26. Ero, R., Leppik, M., Reier, K., Liiv, A. & Remme, J. Ribosomal RNA modification enzymes stimulate large ribosome subunit assembly in E. coli. Nucleic Acids Res. 1, 13–14 (2013).

27. Schwalm, E. L., Grove, T. L., Booker, S. J. & Boal, A. K. Crystallographic capture of a radical S-adenosylmethionine enzyme in the act of modifying tRNA. Science 352, 309–312 (2016).

28. Fitzsimmons, C. M. & Fujimori, D. G. Determinants of tRNA Recognition by the Radical SAM Enzyme RlmN. PloS One 11, e0167298 (2016).

29. Giessing, A. M. B. et al. Identification of 8-methyladenosine as the modification catalyzed by the radical SAM methyltransferase Cfr that confers antibiotic resistance in bacteria. RNA 15, 327–336 (2009).

30. Tsai, K. et al. Directed evolution of the rRNA methylating enzyme Cfr reveals molecular basis of antibiotic resistance. eLife 11, (2022).

31. Aleksandrova, E. V. et al. Structural basis of Cfr-mediated antimicrobial resistance and mechanisms to evade it. Nat. Chem. Biol. 2024 1–10 (2024) doi:10.1038/s41589-023-01525-w.

32. Tsai, K. et al. Structural basis for context-specific inhibition of translation by oxazolidinone antibiotics. Nat. Struct. Mol. Biol. 2022 292 29, 162–171 (2022).

33. Kaminska, K. H. et al. Insights into the structure, function and evolution of the radical-SAM 23S rRNA methyltransferase Cfr that confers antibiotic resistance in bacteria. Nucleic Acids Res. 38, 1652–1663 (2010).

34. Atkinson, G. C. et al. Distinction between the Cfr methyltransferase conferring antibiotic resistance and the housekeeping RlmN methyltransferase. Antimicrob. Agents Chemother. 57, 4019–4026 (2013).

35. Stojkoviæ, V., Noda-Garcia, L., Tawfik, D. S. & Fujimori, D. G. Antibiotic resistance evolved via inactivation of a ribosomal RNA methylating enzyme. Nucleic Acids Res. 44, 8897–8907 (2016).

36. Hansen, L. H. & Vester, B. A cfr-like gene from Clostridium difficile confers multiple antibiotic resistance by the same mechanism as the cfr gene. Antimicrob. Agents Chemother. 59, 5841–5843 (2015).

37. Goldenzweig, A. et al. Automated Structure- and Sequence-Based Design of Proteins for High Bacterial Expression and Stability. Mol. Cell 63, 337–346 (2016).

38. Boltz-2: Towards Accurate and Efficient Binding Affinity Prediction | bioRxiv. https://www.biorxiv.org/content/10.1101/2025.06.14.659707v1.

39. Brilot, A. F., Korostelev, A. A., Ermolenko, D. N. & Grigorieff, N. Structure of the ribosome with elongation factor G trapped in the pretranslocation state. Proc. Natl. Acad. Sci. U. S. A. 110, 20994–20999 (2013).

40. Schaenzer, A. J. et al. Angucyclinones rescue PhLOPSA antibiotic activity by inhibiting Cfr-dependent antibiotic resistance. mBio 14, (2023).

41. Stojkoviæ, V. et al. Cfr(B), cfr(C), and a New cfr-Like Gene, cfr(E), in Clostridium difficile Strains Recovered across Latin America. Antimicrob. Agents Chemother. 64, (2020).

42. van Kempen, M. et al. Fast and accurate protein structure search with Foldseek. Nat. Biotechnol. 1–4 (2023) doi:10.1038/s41587-023-01773-0.

43. Copp, J. N., Anderson, D. W., Akiva, E., Babbitt, P. C. & Tokuriki, N. Exploring the sequence, function, and evolutionary space of protein superfamilies using sequence similarity networks and phylogenetic reconstructions. Methods Enzymol. 620, 315–347 (2019).

44. Cherepanov, P. P. & Wackernagel, W. Gene disruption in Escherichia coli: TcR and KmR cassettes with the option of Flp-catalyzed excision of the antibiotic-resistance determinant. Gene 158, 9–14 (1995).

45. Zheng, W. et al. Folding non-homologous proteins by coupling deep-learning contact maps with I-TASSER assembly simulations. Cell Rep. Methods 1, 100014 (2021).

46. Venkatesan, M. et al. Molecular mechanism of plasmid-borne resistance to sulfonamide antibiotics. Nat. Commun. 14, 1–17 (2023).

47. Minor, W., Cymborowski, M., Otwinowski, Z. & Chruszcz, M. HKL-3000: The integration of data reduction and structure solution - From diffraction images to an initial model in minutes. Acta Crystallogr. D Biol. Crystallogr. 62, 859–866 (2006).

48. Keegan, R. M. & Winn, M. D. Automated search-model discovery and preparation for structure solution by molecular replacement. Acta Crystallogr. D Biol. Crystallogr. 63, 447–457 (2007).

49. Liebschner, D. et al. Macromolecular structure determination using X-rays, neutrons and electrons: recent developments in Phenix. Acta Crystallogr. Sect. Struct. Biol. 75, 861–877 (2019).

50. Emsley, P., Lohkamp, B., Scott, W. G. & Cowtan, K. Features and development of Coot. Acta Crystallogr. D Biol. Crystallogr. 66, 486–501 (2010).

51. Wiegand, I., Hilpert, K. & Hancock, R. E. W. Agar and broth dilution methods to determine the minimal inhibitory concentration (MIC) of antimicrobial substances. Nat. Protoc. 3, 163–175 (2008).

