## Supporting Information for "Structural analysis of 23S rRNA methylating enzyme Cfr reveals RNA binding determinants for methylation regiospecificity and antibiotic resistance"

**Supporting Information Figure S1 - Comparison of 23S rRNA coordinates from the Cfr-rRNA model and crystal structure of *E. coli* ribosome**

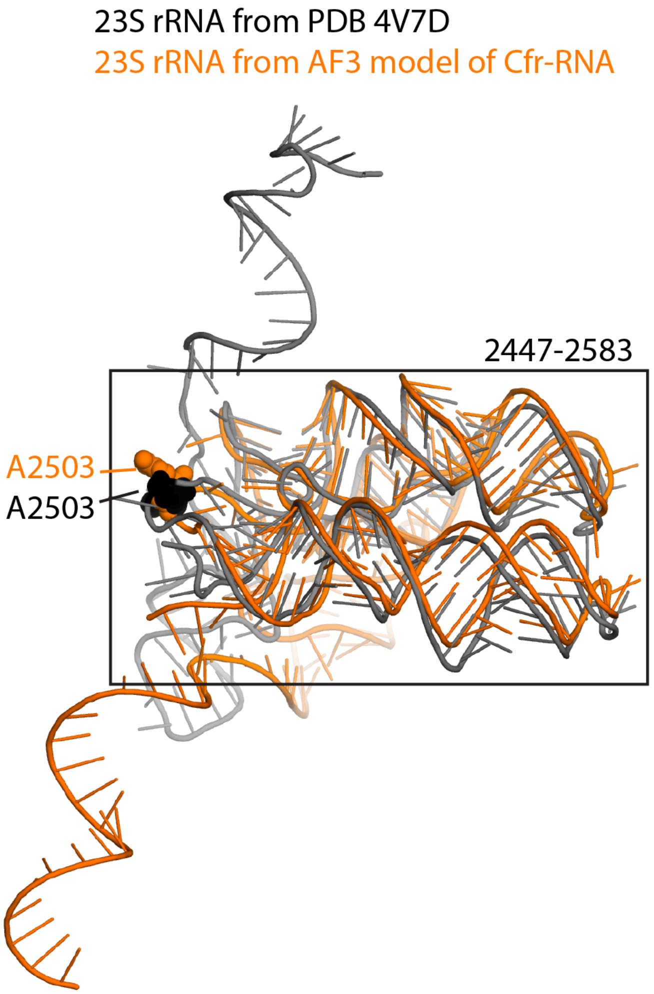


Overlay of coordinates of rRNA used for generation of Cfr-rRNA model (bases 2447-2635) and the equivalent region from crystal structure of the *E. coli* ribosome (PDB 4v7d, doi:10.1073/pnas.1311423110). Comparison shows high structural similarity in 2447-2583 region, which is boxed. A2503 from the model and crystal structure are labelled. AF3, AlphaFold 3.

**Supporting Information Figure S2 - Surface electrostatic charge of Cfr (from Cfr-rRNA AlphaFold3 model)**

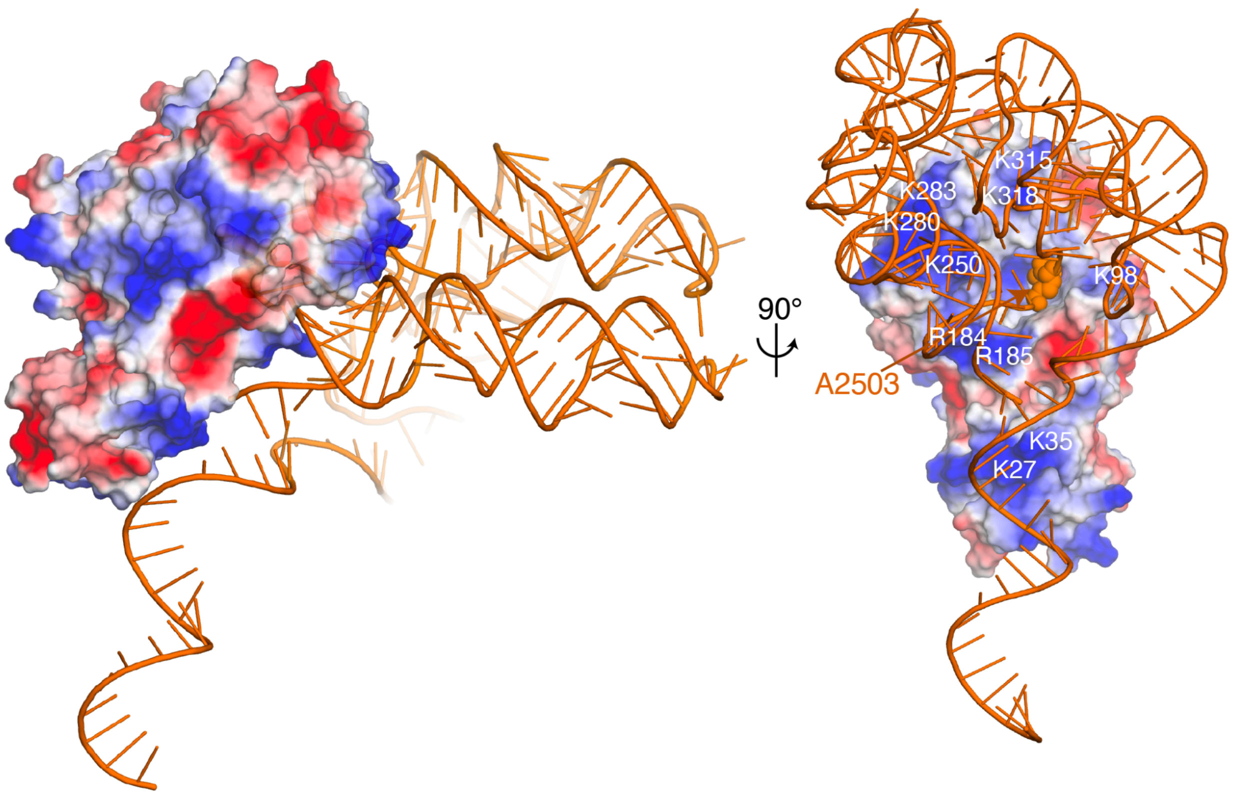


Shown are the coordinates of Cfr and rRNA from the AlphaFold3-generated model. Solvent-accessible surface area of Cfr is shown and colored in a gradient according to electrostatic potential (red = acidic, white = neutral, blue = basic). On the right, basic amino acids that interact with the rRNA are labelled.

**Supporting Information Figure S3 - Evaluation of chloramphenicol resistance conferred by Cfr$^{PROSS-8}$**

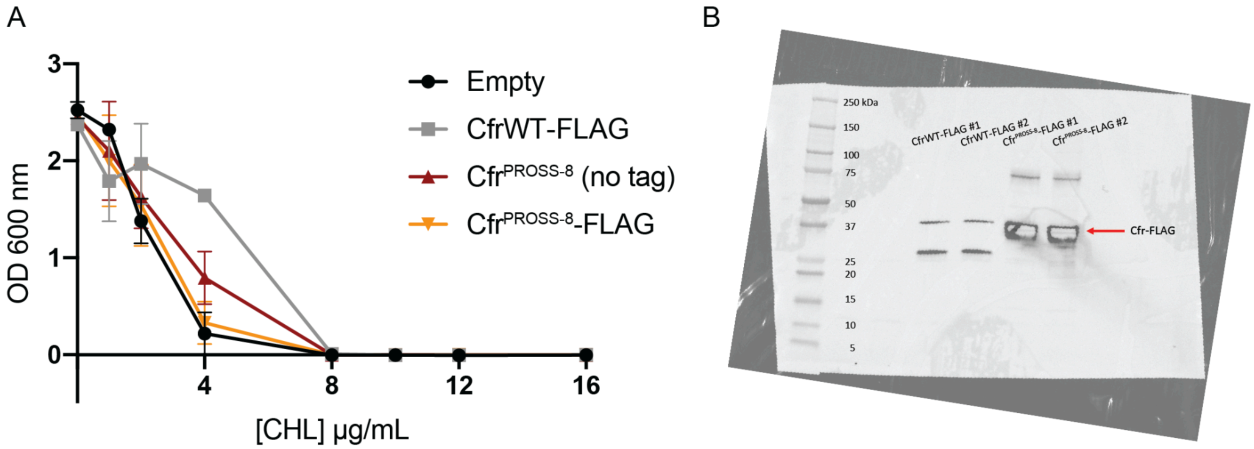


A) CfrWT and Cfr$^{PROSS-8}$ were expressed in *E. coli* BW25113 on the pZA vector with a C-terminal FLAG tag or with no tag, and grown in the presence of different concentrations of chloramphenicol. MIC for CfrWT-FLAG was 8 μg/mL, Cfr$^{PROSS-8}$-FLAG was 4 μg/mL, empty vector control was 4 μg/mL. B) Western blot showing robust expression of CfrWT-FLAG and Cfr$^{PROSS-8}$-FLAG from two replicates each.

**Supporting Information Figure S4 - Validation of expression of Cfr variants in *E. coli BW25113* Δ*bamB*Δ*tolC* and Δ*bamB*Δ*tolC*Δ*rlmN***

A

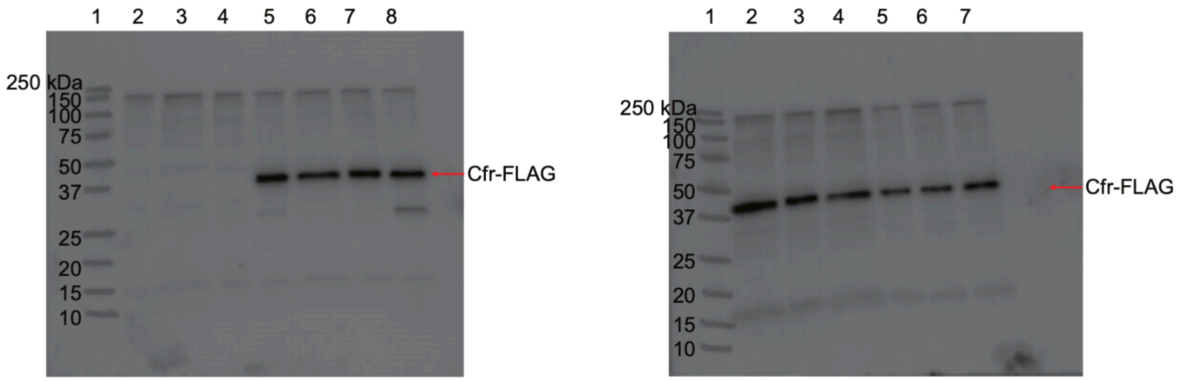


1-MW marker
2-BW25113
3-BW25113 Δ*bamB*Δ*tolC*
4-BW25113 Δ*bamB*Δ*tolC* :: pGDP2 empty vector
5-BW25113 Δ*bamB*Δ*tolC* :: pGDP2-CfrWT
6-BW25113 Δ*bamB*Δ*tolC* :: pGDP2-Cfr M95I
7-BW25113 Δ*bamB*Δ*tolC* :: pGDP2-Cfr Y97E
8-BW25113 Δ*bamB*Δ*tolC* :: pGDP2-Cfr W101R

1-MW marker
2-BW25113 Δ*bamB*Δ*tolC* :: pGDP2-CfrWT
3-BW25113 Δ*bamB*Δ*tolC* :: pGDP2-Cfr S103T
4-BW25113 Δ*bamB*Δ*tolC* :: pGDP2-Cfr S151N
5-BW25113 Δ*bamB*Δ*tolC* :: pGDP2-Cfr S328K
6-BW25113 Δ*bamB*Δ*tolC* :: pGDP2-Cfr Q329T
7-BW25113 Δ*bamB*Δ*tolC* :: pGDP2-Cfr F330R

B

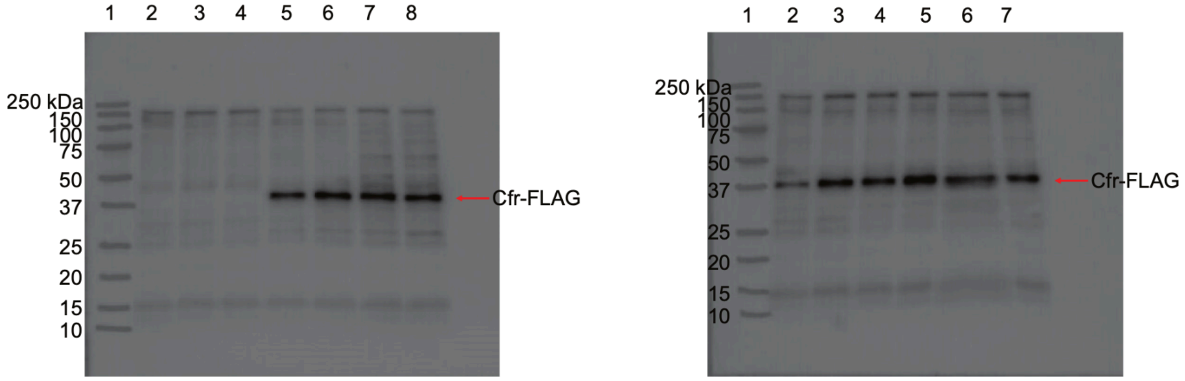


1-MW marker
2-BW25113
3-BW25113 Δ*bamB*Δ*tolC*Δ*rlmN*
4-BW25113 Δ*bamB*Δ*tolC*Δ*rlmN* :: pGDP2 empty vector
5-BW25113 Δ*bamB*Δ*tolC*Δ*rlmN* :: pGDP2-CfrWT
6-BW25113 Δ*bamB*Δ*tolC*Δ*rlmN* :: pGDP2-Cfr M95I
7-BW25113 Δ*bamB*Δ*tolC*Δ*rlmN* :: pGDP2-Cfr Y97E
8-BW25113 Δ*bamB*Δ*tolC*Δ*rlmN* :: pGDP2-Cfr W101R

1-MW marker
2-BW25113 Δ*bamB*Δ*tolC*Δ*rlmN* :: pGDP2-CfrWT
3-BW25113 Δ*bamB*Δ*tolC*Δ*rlmN* :: pGDP2-Cfr S103T
4-BW25113 Δ*bamB*Δ*tolC*Δ*rlmN* :: pGDP2-Cfr S151N
5-BW25113 Δ*bamB*Δ*tolC*Δ*rlmN* :: pGDP2-Cfr S328K
6-BW25113 Δ*bamB*Δ*tolC*Δ*rlmN* :: pGDP2-Cfr Q329T
7-BW25113 Δ*bamB*Δ*tolC*Δ*rlmN* :: pGDP2-Cfr F330R

Western blots of CfrWT and variants expressed from pGDP2 in A) *E. coli* BW25113 Δ*bamB*Δ*tolC* and B) *E. coli* BW25113 Δ*bamB*Δ*tolC*Δ*rlmN*. Each gel is representative of 4 biological replicates.

**Supporting Information Table S1 - X-ray crystallographic statistics for Cfr$^{PROSS-8}$**

| PDB code | 9P0P |
|---|---|
| *Data collection* | |
| Space group | $P2_12_12_1$ |
| Cell dimensions | |
| *a*, *b, c* (Å) | 45.32, 58.61, 115.95 |
| α, β, γ, (°) | 90, 90, 90 |
| Resolution, Å | 50.00 – 2.50 |
| $R_{merge}$[a] | 0.075 (0.882)* |
| $R_{pim}$[b] | 0.027 (0.324) |
| $CC_{1/2}$* | 0.687 |
| $I / \sigma(I)$ | 32.42 (1.6) |
| Completeness, % | 94.1 (72.6) |
| Redundancy | 9.0 (7.3) |
| *Refinement* | |
| Resolution, Å | 42.21 – 2.50 |
| No. unique reflections: working, test | 10508, 925 |
| $R_{work}/R_{free}$[c] | 25.4/30.9 (40.3/48.5) |
| No. atoms | |
| Protein | 2482 |
| Water | 1 |
| *B*-factors | |
| Protein | 84.9 |
| Water | 58.0 |
| R.m.s. deviations | |
| Bond lengths, Å | 0.005 |
| Bond angles, ° | 0.733 |
| Ramachandran plot | |
| Favored, % | 92.3 |
| Allowed, % | 7.7 |
| Outliers, % | 0 |

*All values in brackets and $CC_{1/2}$ values refer to the highest resolution shells.
[a]$R_{merge} = \Sigma_{hkl}\Sigma_j|I_{hkl.j}-I_{hkl}|/\Sigma_{hkl}\Sigma_j I_{hk,j}$, where $I_{hkl,j}$ and $\langle I_{hk,j}\rangle$ are the *j*th and mean measurement of the intensity of reflection *j*.

$^{b}R_{pim} = \Sigma_{hkl}\sqrt{(n/n\text{-}1)}\ \Sigma^{n}_{j=1}|I_{hkl,j}\text{-}I_{hkl}|/\Sigma_{hkl}\Sigma_{j}I_{hk,j}$
$^{c}R = \Sigma|F_p^{obs} - F_p^{calc}|/\Sigma F_p^{obs}$, where $F_p^{obs}$ and $F_p^{calc}$ are the observed and calculated structure factor amplitudes, respectively.
N/A = not applicable.

**Supporting Information Table S2** - *E. coli* strains used in this study

| | Strain | Description | Source or Reference |
|---|---|---|---|
| | *E. coli BW25113* | *E. coli* K-12 BW25113 wild type: Δ(*araD*-*araB*)567 Δ*lacZ*4787(::*rrnB*-3) *rph*-1 Δ(*rhaDrhaB*)568 *hsdR*514 | Dharmacon Inc. |
| 1 | *E. coli* BW25113 Δ*bamB*Δ*tolC* | Hyperpermeable strain | DOI: 10.1016/j.chembiol.2016.11.011 |
| 2 | *E. coli* BW25113 Δ*bamB*Δ*tolC*Δ*rlmN::Kan*$^{R}$ | Hyperpermeable strain; KAN$^{R}$ cassette inserted into *rlmN* locus | DOI: 10.1128/mbio.01791-23 |
| | *E. coli* BW25113 Δ*bamB*Δ*tolC*Δ*rlmN* | Hyperpermeable strain, with the kanamycin cassette inserted into the *rlmN* locus, was eliminated (markerless deletion) | This study |

**Supporting Information Table s3** - Plasmids used in this study

| | Vector | Description | Reference |
|---|---|---|---|
| 1 | pZA-Empty | *E. coli* expression vector Ptet, AMP$^{R}$ | DOI: 10.1093/nar/gkw699 |
| 2 | pZA-Cfr | *E. coli* expression vector Ptet, AMP$^{R}$ encoding *S. aureus cfr* | DOI: 10.7554/eLife.70017 |
| | pCP20 | A temperature-sensitive plasmid expressing FLP recombinase; *Amp*$^{R}$ | DOI: 10.1016/0378 |

| | | | |
|---|---|---|---|
| | | | -1119(95)001 93-a |
| | pGDP2 | *E. coli* gene expression vector, $Km^R$ | DOI: 10.1016/j.chembiol.2016.11.011 |
| | pGDP2::$cfr_{WT}$ | | This study |
| | pGDP2::$cfr_{\mathrm{M95I}}$ | | This study |
| | pGDP2::$cfr_{Y97E}$ | | This study |
| | pGDP2::$cfr_{W101R}$ | | This study |
| | pGDP2::$cfr_{S103T}$ | | This study |
| | pGDP2::$cfrS_{151N}$ | | This study |
| | pGDP2::$cfr_{S328K}$ | | This study |
| | pGDP2::$cfr_{Q329T}$ | | This study |
| | pGDP2::$cfr_{F330R}$ | | This study |

**Supporting Information Table S4 -** oligonucleotides used in this study

| | Oligos | Sequence | Description |
|---|---|---|---|
| 1 | F330R Mutant | 5'-acaattagaagtcaacgtgggattgatatt-3'<br>5'-aatatcaatcccacgttgacttctaattgt-3' | For mutagenesis of pZA vector |
| 2 | W101R Mutant | 5'-aaagcaggtagagagtcattttgt-3'<br>5'-acacccaaaattacatccgcatt-3' | For mutagenesis of pZA vector |
| 3 | S103T Mutant | 5'-gcaggttgggagacattttgtatatca-3'<br>5'-tgatatacaaaatgtctcccaacctgc-3' | For mutagenesis of pZA vector |

| | | | |
|---|---|---|---|
| 4 | S328K Mutant | 5'-acaattagaaaacaatttgggatt-3'<br>5'-aatcccaaattgttttctaattgt-3' | For mutagenesis of pZA vector |
| 5 | Q329T Mutant | 5'-attagaagtacgtttgggattgatatt-3'<br>5'-aatatcaatcccaaacgtacttctaattgt-3' | For mutagenesis of pZA vector |
| 6 | Y97E Mutant | 5'-aacatgaaggagaaagcaggttgg-3'<br>5'-acacccaaaattacatccgcatt-3' | For mutagenesis of pZA vector |
| 7 | S151N Mutant | 5'-catcaaattgataatatttcttttatg-3'<br>5'-cataaaagaaatattatcaatttgatg-3' | For mutagenesis of pZA vector |
| 8 | C2480-C2520 | 5'-gatgtgatgagccgacatcgaggtgccaaacaccgccgtcg-3' | Complementary sequence for rRNA fragment containing A2503 |
